# *Sapap3* knockout mice show threat bias under conflict during platform mediated avoidance task: implications for obsessive compulsive disorder

**DOI:** 10.1101/2024.02.12.579940

**Authors:** Elizabeth E Manning, Elizabeth A Crummy, Zoe LaPalombara, Xiaojun Li, Samyuktha Manikandan, Susanne E Ahmari

## Abstract

Obsessive compulsive disorder (OCD) typically involves cycling between symptoms of intrusive aversive thoughts (obsessions) and repetitive rituals aimed at avoiding these aversive outcomes (compulsions), which interferes with patients’ engagement with other important aspects of their lives. This cycling relationship between obsessions and compulsions highlights a potential role of impaired threat processing and avoidance behavior in OCD symptoms. The most effective behavioral therapy for OCD, exposure with response prevention (ERP), aims to break this cycle. However, it can be difficult for patients to access and engage with, suggesting a need for improved understanding of the neural mechanisms of threat processing and avoidance behavior in OCD to guide development of new and more effective treatments. Platform mediated avoidance (PMA) has proven to be a useful translational paradigm for use in rodents to examine avoidance neurobiology and models relevant to OCD, such as ERP and overtraining-induced persistent avoidance. However, to date this protocol has only been used in rats, and studies in transgenic mouse models relevant to OCD may shed further light on neural mechanisms relevant to disturbances in avoidance and threat processing in the disorder.

To address this gap, we tested *Sapap3* knockout (KO) mice, a leading preclinical model in OCD research, in the PMA task. Using this paradigm, we examined avoidance acquisition, expression, and extinction, as well as reward seeking under motivational conflict in two separate cohorts conditioned using higher (0.4 mA) or lower (0.24 mA) intensity shock. Surprisingly, the most striking difference observed in *Sapap3*-KOs vs control mice was heightened suppression of lever pressing for rewards during a tone signaling impending threat, suggesting a shift in action selection under motivational conflict (genotype effect avoidance conditioning: 0.24 mA cohort p=0.011, 0.4 mA cohort p=0.07; avoidance extinction: 0.24 mA p=0.057, 0.4 mA p=0.042). Avoidance responding was also acquired more slowly in *Sapap3*-KOs trained with a low intensity shock (time x genotype interaction p=0.025) and was extinguished more robustly following ERP (genotype effect p=0.043). In contrast, avoidance was similar between *Sapap*3 KOs and WT littermate controls trained using higher intensity shock (0.4 mA cohort time effect p<0.0001). Expression of the immediate early gene c-Fos associated with reinstatement of avoidance after ERP showed preliminary evidence for decreased activity of medial orbitofrontal cortex (mOFC) in KOs which may contribute to observed differences in PMA performance (KO vs WT mOFC c-Fos t-test p=0.0456). Together these findings suggest that mOFC dysfunction may contribute to increases in the influence of threats vs rewards over action selection in an animal model with relevance to OCD, and that the *Sapap3*-KO model presents valuable opportunities for deeper mechanistic investigation of avoidance and threat processing relevant to the human disorder.

## Introduction

Obsessive-compulsive disorder (OCD) is a chronic, debilitating disorder affecting 1-3% of the population. It is characterized by recurring distressing thoughts (obsessions) and ritualized actions (i.e. compulsions) that are often performed to avoid the perceived harmful outcomes of obsessions and obtain relief from obsession-related distress (Pauls et al., 2014). The link between obsessions, compulsions, and avoidance is further highlighted by studies showing that impairments in threat processing and avoidance are associated with compulsion severity and poorer treatment outcomes in OCD (McGuire et al., 2012; Wheaton et al., 2018; Wheaton et al., 2021). Additionally, OCD patients exhibit biases to attend to and learn to avoid negative stimuli (Endrass et al., 2011; Kajs et al., 2022), habitually avoid devalued threats (Gillan et al., 2014), have devaluation impairments on a rule-based avoidance task with longer illness duration (Chase et al., 2020), and inappropriately perform avoidant behaviors to safety cues (De Kleine et al., 2023). Together, these studies suggest that active avoidance is a persistent component that could contribute to the development and severity of OCD, but little is known about the mechanisms underlying its dysregulation.

Active avoidance is a defensive behavioral response that is commonly modeled as performing an action to prevent exposure to a threat. Common preclinical models include shuttle avoidance, in which rodents are conditioned to run across a chamber to avoid shocks administered on one half of a shuttlebox, as well as platform avoidance, in which rodents can avoid shocks by stepping onto a platform to prevent contact with a grid floor delivering a foot shock. The platform-mediated avoidance task (PMA) specifically involves rodents choosing between lever pressing for rewards or making an avoidance response by mounting a platform during a cue that predicts a foot shock, thereby modeling conflict-based active avoidance, a critical feature of maladaptive avoidant behaviors (Ball & Gunaydin, 2022; Diehl et al., 2019). Extent of PMA training impacts avoidance extinction in models of exposure with response prevention (ERP) (Martinez-Rivera et al., 2020), a well-established treatment for OCD (Abramowitz, 2006), making this a powerful translational model for understanding the development of and alterations to avoidance behaviors in OCD (Rodriguez-Romaguera et al., 2016). Studies in rodents have identified circuits involved in PMA conditioning and extinction whose homologues are also implicated in neuroimaging studies of avoidance in OCD patients, including medial prefrontal cortex and striatal regions relevant to threat appraisal and motivated behaviors altered in OCD (Diehl et al., 2018; Gillan et al., 2015; Martinez-Rivera et al., 2023; Rodriguez-Romaguera et al., 2016). Despite this progress in using PMA to understand the neurobiology of avoidance relevant to OCD, to date it has only been used in behavioral models relevant to OCD (e.g. overtraining, ERP). Testing of OCD-relevant transgenic mice on this task may therefore provide important opportunities for identifying disease relevant mechanisms.

*Sapap3* knockout mice (KOs) are a common model for OCD-related behaviors, exhibiting compulsive grooming, elevated anxiety (Lamothe et al., 2023; van den Boom et al., 2019; Welch et al., 2007), and deficits in behavioral flexibility (Hadjas et al., 2019; Manning et al., 2019; van den Boom et al., 2019) which map onto symptoms observed in clinical populations (Benzina et al., 2021; Burguiere et al., 2015). Recent studies also demonstrate enhanced fear conditioning and impaired fear extinction in KOs [(Kajs et al., 2022), LaPalombara et al., unpublished observations], suggesting heightened sensitivity to negative stimuli and threat appraisal. However, KO performance in active avoidance tasks has not been investigated. Identifying behavioral and neurobiological changes associated with avoidance acquisition, extinction, or conflict-based reward seeking in *Sapap3*-KOs using the PMA task could prove valuable for understanding alterations in avoidance behavior and associated symptoms in OCD.

To address this gap, we examined PMA acquisition and extinction using an ERP model in male *Sapap3*-KOs and wild-type (WT) littermate controls, followed by analysis of the immediate early gene c-Fos to begin to examine potential neural mechanisms involved in reinstatement of active avoidance following ERP. We hypothesized that KOs would show excessive avoidance responding, even following ERP exposure, similar to effects previously observed with overtraining or orbitofrontal cortex disruptions (Martinez-Rivera et al., 2020; Rodriguez-Romaguera et al., 2016).

## Methods

### Animals

Male *Sapap3*-KOs and WT littermates were generated through heterozygous x heterozygous breeding, and maintained on C57BL/6 background. Mice were group-housed with 2–4 mice per cage, with *ad libitum* access to food and water until operant training commenced at approximately 4 months of age, when they subsequently had restricted daily food access and *ad libitum* access to water. Mice were housed in a reverse light-cycle room starting at least 1 week prior to the start of operant testing (12:12, lights on at 7:00 pm). All procedures were carried out in accordance with the guidelines for the care and use of laboratory animals from the National Institutes of Health (NIH) and with approval from the University of Pittsburgh Institutional Animal Care and Use Committee (IACUC).

### Behavior testing

#### Grooming test

Approximately one week prior to the beginning of operant training when animals were still housed in regular light cycle, mice were tested in acrylic testing chambers (8 × 8 × 12 inches) during their light cycle similar to as previously described (Manning, Wang, et al., 2021). Videos were recorded using a front-on view of the testing arena and scored offline for time spent grooming. Mice were an average of 3.7 months of age (range 3.1-4.1) at the time of the grooming test.

### Operant acquisition

Throughout operant training, mice were food restricted to 85-90% of their free feeding weight, with restricted access to food for 1-4 hours daily following their daily operant training. During daily feeding, mice were single housed and given 1-3g food (regular chow), with the amount given adjusted for their individual body weight to maintain the target range.

Mice were on average 4.1 months of age (range 3.5-4.5) at the time when operant training commenced. First, mice were habituated to the food reward used during operant training (20-mg chocolate-flavored grain-based pellets; BioServ, Flemington, NJ) in their home cage on two separate occasions to prevent neophobia. Next, mice were habituated to the operant chamber (MED-307W, Med Associates, Fairfax, VT) for 2-3 days, with the fan on and house light on covered by a red filter, and access to 10 free pellets in the food magazine. Mice then progressed to training on a fixed ratio 1 (FR1) schedule, where one lever was available and every lever press resulted in delivery of a food reward, with a maximum of 20 rewards available in each 1 hour session. Once mice reached 20 rewards within the hour (typically 1-5 days), they progressed to 30 minute duration variable ratio 2 (VR2) sessions, where 1, 2 or 3 presses were required before a reward was delivered. Once mice achieved a criteria of >30 pellets in a session on 3 consecutive days, they moved on to variable interval 30 (VI-30) sessions, where rewards were delivered on the first press after a variable interval that was an average of 30 seconds long. At this stage of training, an acrylic platform was introduced to the chamber in the corner opposite the lever, to allow mice to habituate to it prior to avoidance conditioning. Mice progressed onto platform avoidance conditioning after at least 2 weeks training on this schedule.

### Platform avoidance conditioning

Following acquisition of high-rate lever pressing on a VI-30 schedule, mice underwent 14 days of platform avoidance conditioning similar to the paradigm previously described in rats (Bravo-Rivera et al., 2014). During daily conditioning, mice were trained in 36 minute long sessions, during which 9 tones (30 seconds duration, average inter-tone interval 3 minutes, first tone starting at 5 minutes 12 seconds) co-terminated in a 2 second foot shock. Mice had access to the same lever on VI-30 schedule throughout testing, and the platform was located in the opposite corner of the chamber. Movement onto the platform at the end of the tone period could be used to avoid the aversive foot shock. One cohort of animals was tested using a foot shock intensity of 0.40mA, and a second cohort was tested using a milder foot shock intensity of 0.24 mA (60% of the original intensity).

Lever press suppression during the tone was calculated using the press rate in the 60 seconds prior to the tone (pretone press rate):

Suppression of lever pressing = (pretone press rate – tone press rate)/(pretone press rate + tone press rate) x100

Video recordings (Genious Wide Cam F-100) of the sessions from above the operant chamber were used to quantify time spent avoiding on the platform. Time spent on the platform was quantified automatically using Anymaze software (Wood Dale, IL), with zones set up for the “platform” and “whole chamber”.

### Platform avoidance extinction with response prevention

Following 14 days of platform avoidance training, mice were trained for 3 days in an extinction with response prevention paradigm, in which access to the platform was blocked by a Perspex barrier (Rodriguez-Romaguera et al., 2016). On each of these 3 days, mice were exposed to 15 tones (average 3 second ITI starting 5 minutes 12 seconds into the session), and no foot shock outcome was delivered. Mice were able to lever press on VI-30 schedule as per conditioning sessions.

### Platform avoidance extinction test

Following extinction with response prevention, mice were tested in a final extinction test session, in which access to the platform was no longer blocked. Similar to conditioning, mice were exposed to 9 tones; however, these no longer co-terminated with a foot shock.

### Immediate early gene analysis: tissue collection and processing

Mice from the 0.24 mA cohort were deeply anesthetized with ketamine 60 minutes after the first tone in the extinction test [as per previous PMA studies (Bravo-Rivera et al., 2015)], and transcardially perfused with 4% paraformaldehyde (PFA; Sigma Aldrich, St. Louis, MO) in 0.1 M phosphate-buffered saline (PBS). Brains were post-fixed in PFA overnight and subsequently transferred to a 30% sucrose solution with 0.1% sodium azide (Sigma) for a minimum of 48 hours. Brains were frozen on dry ice and sliced in 35µm sections on a Leica cryostat. Free-floating sections were stored in PBS with 0.1% sodium azide at 4°C until used for immunohistochemistry.

### Immunohistochemistry

Similar to previous studies (Manning et al., 2019), tissue was washed 3x with 0.1 M tris-buffered saline (TBS; Sigma) prior to incubating in 1% hydrogen peroxide for 10 minutes. After additional TBS washing, sections were incubated in a blocking solution (3% normal goat serum in TBS) for 30 minutes before incubating in c-Fos primary antibody for 2 days (rabbit anti c-Fos, 1:1000; Millipore, Burlington, MA). Sections were then washed in 3x in TBS with 0.3% Triton X-100 (Sigma) for 10 minutes prior to incubating in a goat anti-rabbit biotinylated secondary antibody (1:500; Vector, Newark, CA) for 2 hours. Sections were then washed in TBS+Triton 3 times for 10 minutes before blocking in a tertiary ABC solution (1:100 in TBS+Triton; Vector ABC Kit) for 1 hour. Sections were washed in TBS 3 times for 10 minutes before being transferred to a 3,3′-Diaminobenzidine chromogen solution (DAB 1:50 in buffer stock solution and H2O2, Sigma) for 5 minutes and subsequently washed 3 times in TBS. Tissue was mounted on charged slides and dried prior to ethanol washes (70%, 80%, 95%, 100% Ethanol, 3 minutes each) and washed in a histological clearing agent (Xylene substitute; Sigma Aldrich; 3 minutes), and then cover slipped with DPX mounting agent (Sigma Aldrich).

### Microscopy and analysis

Slides were imaged at 20x using a slide-scanning scope (VS120, Olympus). c-Fos was automatically detected and quantified using CellSense. c-Fos density was calculated using the average c-Fos count within a 300 µm x 300 µm area in each hemisphere for 2-3 sections per animal.

### Statistical analysis

As KOs are well-documented to have increased grooming, a one-tailed t-test was used to detect genotype differences in time spent grooming. Lever suppression was calculated across the session or within three-trial bins (ERP and extinction test). Two-way repeated measures ANOVA or restricted maximum likelihood (REML) with Geisser-Greenhouse corrections for sphericity (ε) were used to detect differences in press rate in VI-30 training and platform mediated avoidance conditioning (i.e. lever suppression, percent of time spent on the platform), ERP lever suppression, and extinction tests of platform reinstatement by genotype or session. Genotype differences in overall lever suppression and lever suppression following the first tone during the extinction test session were analysed with unpaired t-tests. Differences in c-Fos density across regions of interest by genotype were analysed with a two-way ANOVA. Bonferroni post hoc tests for multiple comparisons were performed when significant interactions were detected. Pearson’s correlations were performed for regional c-Fos in each ROI to determine if there were associations between c-Fos expression and lever suppression to the first tone in the extinction test. 4 mice were excluded from analyses (0.4 mA cohort: 2 KOs; 0.24 mA cohort: 1 WT and 1 KO) that did not acquire avoidance in 14 days of conditioning. One subject (KO) was excluded from c-Fos analyses in mOFC because its value was a statistical outlier (>2 standard deviations above the mean). Graphpad Prism 10 was used for all analyses.

## Results

*Sapap3*-KOs displayed a compulsive grooming phenotype, exhibiting significantly longer time spent grooming than WT littermate controls (Figure 1B one-tailed t-test t(22) =3.5, p = 0.0011). Mice were trained to lever press for reward pellets, acquiring significantly higher press rates over a two-week training period at VI-30 with similar performance between genotypes (Supplementary Figure S1A: two-way RM ANOVA 0.4 mA cohort: main effect of time F(3.428, 34.28) = 5.2, ε = 0.21, p = 0.0035, main effect of genotype F(1, 10) = 0.03, ns; interaction of time x genotype F(16, 160) = 3.9, p<0.0001; S1B: REML 0.24 mA cohort: main effect of time F (2.62, 30.11) = 3.7, ε =0.15, p= 0.026, main effect of genotype F (1, 12) = 0.3489, ns, interaction of time x genotype F (18, 207) = 0.3, ns).

**Figure 1.**
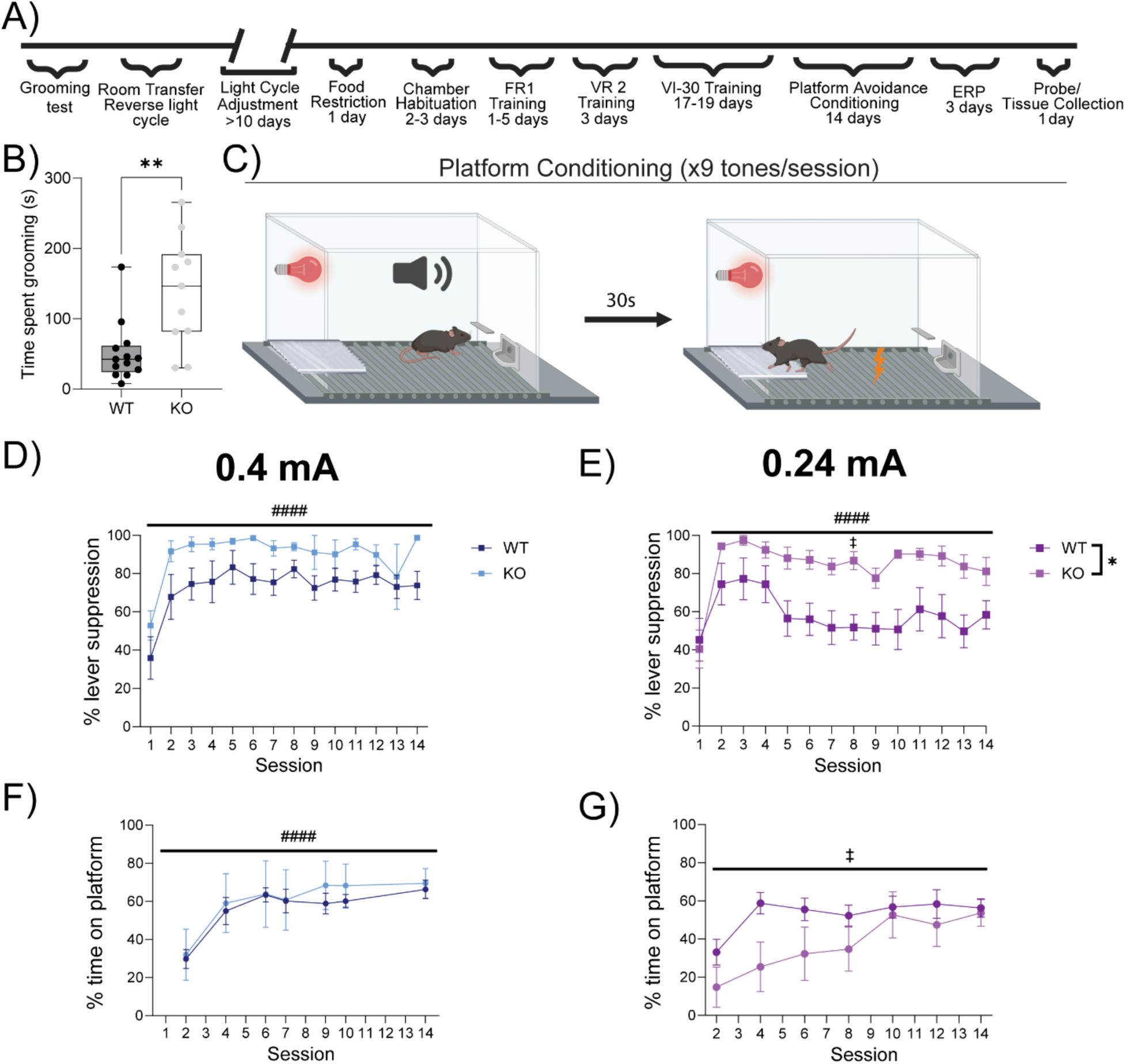
*Sapap3*-KOs are more sensitive to reward seeking under threat conditions. A) Timeline of reward training, platform avoidance, extinction-response prevention, and extinction test of platform avoidance. B) KOs groom more than WTs at ∼4 months of age. C) Schematic of platform-mediated avoidance sessions, where mice can press a lever to obtain rewards on a VI30 schedule, but at risk of foot shock following a warning tone. Mice can successfully avoid shocks by entering onto a platform, preventing contact with the shock floor. D) Platform avoidance conditioning with a 0.4 mA foot shock results in high levels of lever suppression with no differences in genotype. E) Milder 0.24 mA foot shock administered during platform avoidance conditioning produces less lever suppression in WT mice than KOs, that increases in both genotypes throughout conditioning. F) Platform avoidance in the 0.4 mA cohort is acquired to similar performance levels regardless of genotype. G) KOs are slower to acquire platform avoidance in early conditioning with 0.24 mA foot shocks than WTs. Grooming: WTs (n=13), KOs (n= 11). 0.4 mA cohort: WTs (n=7), KOs (n=5); 0.24 mA cohort: WTs (n=7), KOs (n=7). #-main effect of time, *-main effect of genotype, ‡ - interaction of time x genotype. #,*,‡-p<0.05; #### -p<0.0001.

Following reward acquisition, mice were trained to enter a platform during a 30 second tone period to avoid either a 0.4 mA or 0.24 mA foot shock that co-terminated with the tone. During platform avoidance, rewards remained accessible throughout the session; however, rewards could not be accessed while the mouse was on the platform, producing a motivational conflict. When mice were trained using a 0.4 mA intensity foot shock, reduced reward-seeking during the aversive tone (i.e. lever suppression) occurred in both KOs and WTs throughout conditioning, with a trend towards greater lever suppression in KOs (Figure 1D: two-way RM ANOVA main effect of time F(3.537, 35.37) = 9.5, ε =0.27, p<0.0001, main effect of genotype F(1, 10) = 4.1, p=0.0703, interaction of time x genotype F(13, 130) = 0.6, ns). No differences were observed between groups in acquisition of platform avoidance across the conditioning period, as measured by time spent on the platform (Figure 1F: REML main effect of time F(2.808, 26.67) = 18.0, ε = 0.23, p<0.0001, main effect of genotype F(1, 10) = 0.001, ns, interaction of time x genotype F(12, 114) = 0.46, ns).

Lower-intensity foot shocks resulted in more pronounced genotype differences in lever suppression across conditioning, with WTs pressing significantly more often for rewards than KOs during the tone period (i.e. showing riskier behavior; Figure 1E: two-way RM ANOVA main effect of time F(4.154, 49.85) = 9.4, ε = 0.32, p<0.0001, main effect of genotype F(1, 12) = 9.0, p=0.011, interaction of time x genotype F(13, 156) = 2.5, p=0.0048; Bonferroni post hoc WT vs. KO session 8 t(10.76) = 4.3, p=0.02). In addition, time spent on the platform was higher in WTs during early training, though both groups acquired platform avoidance and spent more time on the platform in later sessions (Figure 1G: two-way RM ANOVA main effect of time F(3.909, 46.91) = 10.4, ε =0.6515, p<0.0001, main effect of genotype F(1, 12) = 1.9, ns, interaction of time x genotype F(6,72) = 2.6, p=0.025); however, no significant post hoc interactions were detected for specific conditioning sessions (Bonferroni post hoc WT vs. KO, ns). When comparing the effect of shock intensity on lever suppression within each genotype, this showed that while lever suppression was decreased in WTs between 0.4 mA and 0.24 mA cohorts, lever suppression was not different in KOs trained with different intensity shock (Supplementary figure S1C (WT): two-way RM ANOVA main effect of time F(4.064, 48.77) = 5.5, ε = 0.31, p-0.001, main effect of shock intensity F (1, 12) = 2.008, ns, interaction of time x shock intensity F(13, 156) = 3.43, p=0.0001, Bonferroni post hoc 0.4-0.24 day 8 t(10.61) = 3.7, p=0.049; S1D (KOs): two-way RM ANOVA: main effect of time F(3.261, 32.61) = 12.3, ε = 0.25, p<0.0001, main effect of shock intensity F(1, 10) = 1.3, ns, interaction of time x shock intensity F(13, 130) = 1.0, ns).

After acquiring platform avoidance, mice underwent three sessions of ERP, in which the shock-paired tones were presented without subsequent shocks, but mice were prevented from accessing the platform and making an avoidance response by a plexiglass barrier. *Sapap3*-KOs exhibited deficits in extinction learning, maintaining higher lever suppression than WTs (Figure 2B: 0.4 mA cohort: REML main effect of time F(4.169, 41.39) = 6.5, ε= 0.2978, p = 0.0003, main effect of genotype F(1, 10) = 5.5, p=0.042, interaction of time x genotype F(14, 139) = 0.9, ns). With lower intensity foot shock conditioning, lever suppression reduction was similar between genotypes, though KOs trended towards greater suppression (Figure 2C 0.24 mA: two-way RM ANOVA main effect of time F(5.779, 69.34) = 3.1, ε=0.4128, p=0.01, main effect of genotype F(1, 12) = 4.4, p=0.057, interaction of time x genotype F (14, 168) = 1.299, ns).

**Figure 2:**
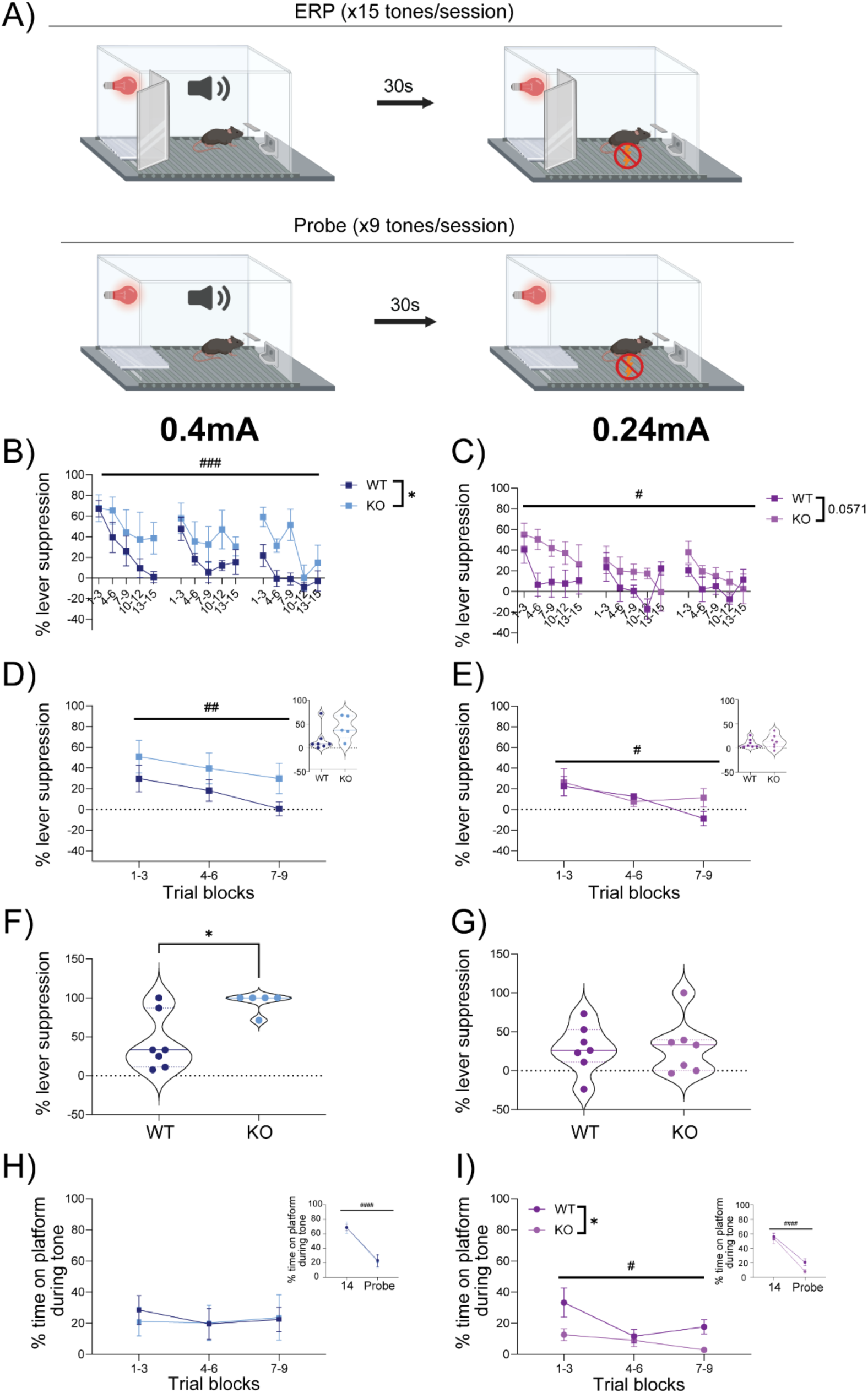
*Sapap3-*KOs maintain diminished reward seeking under extinction, but extinguish platform avoidance similarly to controls. A) Schematic of extinction-response prevention sessions (top) with a barrier to prevent platform avoidance during tone presentations, and an extinction test session (bottom) where access to the platform is reinstated. B) Following 0.4 mA foot shock avoidance conditioning, WTs and KOs increasingly engaged in reward seeking (reduced suppression) across ERP, but KOs maintained significantly higher lever suppression. C) Following 0.24 mA foot shock avoidance conditioning, WTs and KOs, increasingly sought rewards, with trends towards greater lever suppression in KOs. D) 0.4 mA cohort lever suppression continued to decrease in WTs and KOs in the extinction test, with no differences in average lever suppression during the session by genotype (inset). E) 0.24 mA cohort lever suppression continued to decrease across the extinction test session with no differences in overall suppression by genotype (inset). F) 0.4 mA cohort lever suppression following the first tone in the extinction test session was significantly higher for KOs than WTs. G) No group differences in lever suppression following the first tone were detected in the 0.24 mA cohort. H) Platform avoidance did not change across the extinction test session and was similar between groups, with decreased overall platform avoidance following ERP versus the final 0.4 mA conditioning session in both KOs and WTs (inset). I) Avoidance decreased throughout the extinction test in the 0.24 mA cohort, and KOs spent less time avoiding on the platform following ERP. However, overall platform avoidance significantly decreased from the final 0.24 mA conditioning session in both KOs and WTs (inset). 0.4 mA cohort: WT (n=7), KO (n=5); 0.24 mA cohort: WT (n=7), KO (n=7). # - main effect of time, * - main effect of genotype. #,* p<0.05, ## p<0.01, ### p<0.001,#### p<0.0001.

Finally, mice were tested in an extinction test session following ERP, in which the platform was available, but no foot shocks were delivered. During this session, reward-seeking was similar between groups in both cohorts and increased throughout the session (Figure 2D: two-way RM ANOVA 0.4 mA main effect of time F(1.888, 18.88) = 4.9, ε= 0.94, p=0.021, main effect of genotype F(1, 10) = 2.6, ns, interaction of time x genotype F(2, 20) = 0.2, ns; inset average across 9 tones: two-tailed t-test t (10)=1.8, ns; Figure 2E: two-way RM ANOVA 0.24 mA main effect of time F(1.844, 22.13) = 4.5, p=0.025, main effect of genotype F(1, 12) = 0.6, ns, interaction of time x genotype F(2, 24) = 1.3, ns; inset average across 9 tones: two-tail t-test t(12)=0.6, ns). When comparing the effect of shock intensity on lever suppression across ERP within each genotype, similar extinction of lever suppression occurred at both shock intensities in both genotypes (Supplementary figure S2A,B: two-way RM ANOVA A) main effect of time F (6.113, 73.36) = 6.326, ε= 0.44, p<0.0001, main effect of shock intensity F (1, 12) = 1.0, ns, interaction of time x shock intensity F (14, 168) = 1.4, ns; REML B) main effect of time F (5.558, 55.19) = 3.8, ε= 0.40, p=0.0038, main effect of shock intensity F (1, 10) = 2.1, ns, interaction of time x shock intensity F (14, 139) = 0.5, ns).

When examining suppression following the first tone of the extinction test, when no additional extinction learning had yet occurred, KOs had significantly more lever suppression than WTs with 0.4 mA conditioning (Figure 2F unpaired t-test, t(10) = 3.0, p=0.01), but no differences in lever suppression at the first tone when conditioned with milder foot shocks (Figure 2G unpaired t-test, t(12)=0.1, ns). Interesting, shock intensity impacted lever suppression to the first tone in KOs only, with elevated suppression in mice tested with high intensity shock, whereas WTs showed no differences between the two cohorts (Supplementary figure 2C,D: unpaired t-test WTs C) t(11) = 0.7, ns; KOs D) t(10) = 3.8, p=0.0035).

Platform avoidance in the extinction test session did not continue to extinguish within session in the 0.4 mA cohort and was similar in both WTs and KOs (Figure 2H two-way RM ANOVA: 0.4 mA main effect of time F(1.467, 14.67) = 0.2, ε=0.74, ns, main effect of genotype F(1, 10)

= 0.02, ns, interaction of time x genotype F(2, 20) = 0.2, ns). In contrast, platform avoidance continued to extinguish during the extinction test in the 0.24 mA cohort, with KOs exhibiting lower overall avoidance than the WTs (Figure 2I two-way RM ANOVA: main effect of time F(1.304, 15.65) = 6.4, ε= 0.65, p=0.0169, main effect of genotype F(1, 12) = 5.1, p=0.043, interaction of time x genotype F(2, 24) = 2.5, ns). In both cohorts, platform avoidance was extinguished following ERP when comparing day 14 of conditioning to the extinction test session, with no significant differences between WT and KOs in platform time (Figure 2H inset: two-way RM ANOVA 0.4 mA main effect of time F(1, 10) = 92.0, p<0.0001, main effect of genotype F(1, 10) = 0.01, ns, interaction of time x genotype F(1, 10) = 0.03, ns; 2G inset: 0.24 mA: main effect of time F(1, 12) = 90.2, p<0.0001, main effect of genotype F(1, 12) = 1.7, ns, interaction of time x genotype F(1, 12) = 1.4, ns). Extinction of platform avoidance was not different between cohorts conditioned with high vs low shock intensity in either of the genotypes, with both cohorts showing strong extinction comparing platform avoidance at the end of conditioning to the extinction session (Supplementary figure S2E,F: two-way RM ANOVA WTs E) main effect of time F (1, 12) = 91.0, p<0.0001, main effect of shock intensity F(1, 12) = 1.1, ns, interaction of time x shock intensity F(1,12) = 1.4, ns; F): main effect of time F(1, 10) = 90.96, p<0.0001, main effect of shock intensity F(1, 10) = 3.37, ns, interaction of time x shock intensity F(1,10)=0.027, ns).

To identify whether any cortical or striatal regions that are relevant to OCD pathophysiology and avoidance learning were differentially activated in KOs during platform avoidance, tissue from the 0.24 mA conditioning cohort was collected to quantify expression of the immediate early gene c-Fos (an indirect static measure of neuronal activation) associated with exposure to the first tone during the extinction test. There were region specific differences between genotypes in c-Fos density following PMA extinction test [Figure 3A restricted maximum likelihood (REML) main effect of region F(2.680, 29.04) = 21.5, p<0.0001, main effect of genotype F(1, 11) = 1.2, ns, interaction of region x genotype F(6, 65) = 2.8, p=0.018]. Although post hoc tests were not significant, exploratory analysis with a two-tailed t-test showed that this interaction was primarily driven by decreased c-Fos expression in the medial orbitofrontal cortex (mOFC) of *Sapap3*-KOs (Figure 3B: t(10)=2.3, p=0.0457). Further exploratory analysis of correlations between lever press suppression during the first tone of the extinction test session and c-Fos in individual ROIs across both genotypes identified a significant association between mOFC c-Fos density and lever suppression (Pearson R=0.586, p=0.045) and a trend towards associations between infralimbic cortex (IL) c-Fos and lever suppression (Pearson R=0.545, p=0.054).

**Figure 3:**
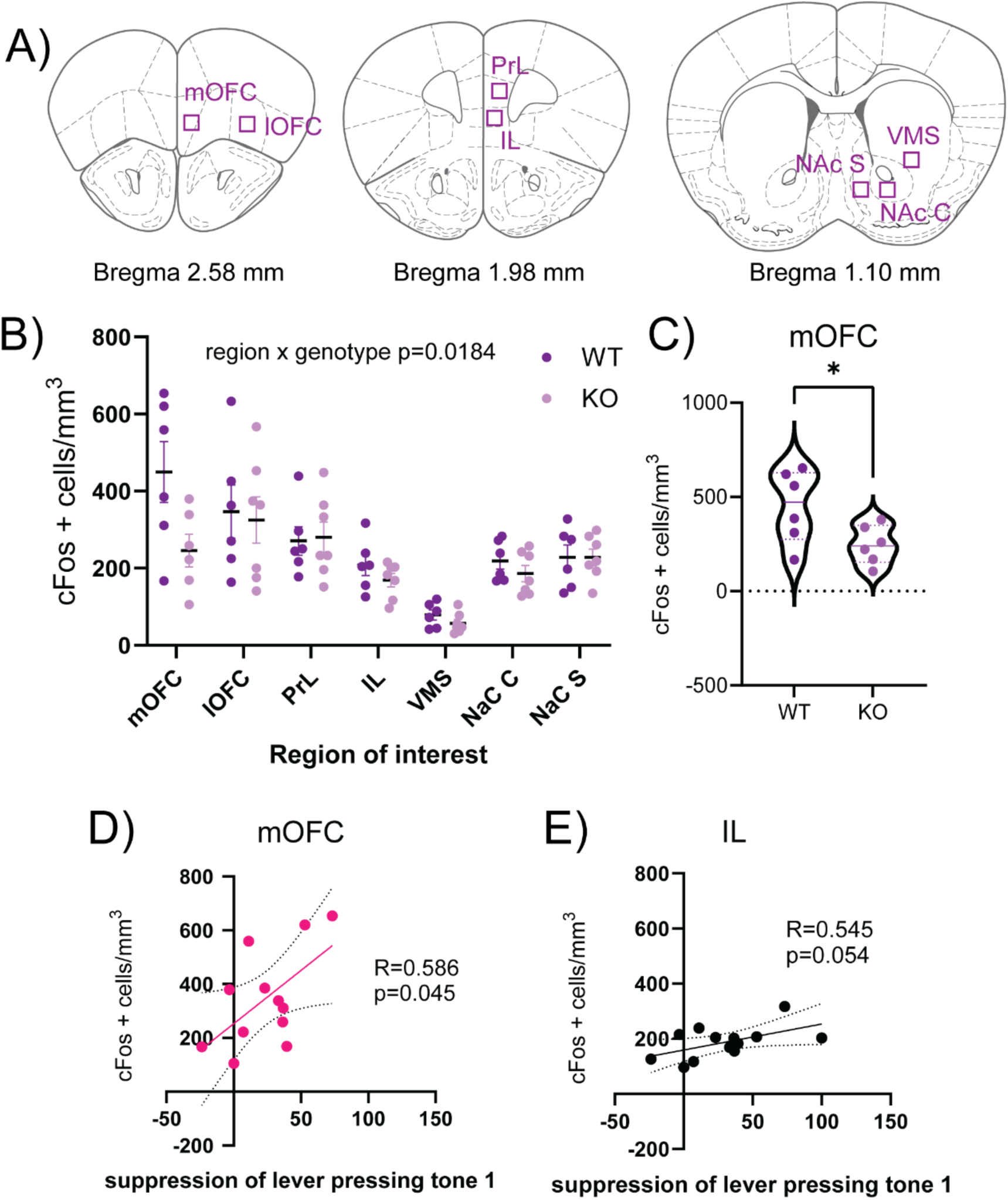
*Sapap3*-KOs show altered neural response to avoidance reinstatement as measured by the immediate early gene c-Fos. A) Schematic of regions of interest in prefrontal cortex and striatum used for c-Fos analysis. B) The density of c-Fos positive cells showed region specific genotype differences. C) Although post hoc tests did not find significant effects in individual regions, the region x genotype interaction was largely driven by decreased c-Fos expression in the mOFC of *Sapap3-*KOs. Exploratory examination of correlations between regional c-Fos density and behavior found that suppression of lever pressing during the first tone of the extinction test session was correlated with c-Fos expression in the mOFC across genotypes (D) and showed a trend for correlation with IL c-Fos (E). WT n=6, KO n=7 (except mOFC where n=6), panels D and E show both genotypes together. * p<0.05

## Discussion

The current study adapted the PMA task for use in mice to examine: 1) changes in avoidance behavior and threat processing, and 2) impact of these changes on reward seeking behavior during motivational conflict, in the *Sapap3*-KO OCD-relevant mouse model. In addition, we used expression of the immediate early gene c-Fos to identify candidate neural structures that may contribute to altered behavior in PMA. We found robust increases in suppression of reward seeking in KOs during the shock-predictive cue, suggesting increased sensitivity of threat processing systems. There was also conflicting evidence for altered extinction in KOs. While KOs tested with a higher intensity shock (0.4 mA) showed elevated suppression of reward-seeking during the first tone of the extinction test session (suggestive of impaired extinction), KOs tested with a lower intensity shock (0.24 mA) showed reduced avoidance in the extinction test session, suggesting enhanced avoidance extinction. We also provide early evidence for a role of altered mOFC activity in changes in threat processing and avoidance in *Sapap3*-KOs through c-Fos expression, which warrants further investigation *in vivo*.

OCD patients are biased towards appraising negative emotional stimuli (Kajs et al., 2022), have faster acquisition from negative versus positive outcomes (Endrass et al., 2011), and show more intolerance and distress during uncertainty, even in situations with low uncertainty or known risk (Jacoby et al., 2023). We therefore wanted to examine avoidance learning in an OCD-relevant mouse model to begin to address mechanisms underlying threat- and risk-assessment. We found that platform avoidance acquisition is similar between WT and *Sapap3*-KOs, with both genotypes acquiring platform avoidance responses within two weeks of training. This suggests that constitutive deletion of the *Sapap3* gene does not impact active avoidance learning, although delayed acquisition of avoidance in mice trained with low intensity footshock (0.24 mA) suggests that there may be some subtle differences between genotypes. However, in addition to examining changes in avoidance learning, the PMA task also assesses reward seeking under motivational conflict via suppression of lever pressing during the shock-paired tone. Our results clearly demonstrate that KOs rapidly reached near-complete suppression of lever pressing during the aversive tone in both cohorts, regardless of the threat magnitude (Supplementary figure 1). In contrast, as would be expected for optimal task performance, lever press suppression was flexibly adjusted based on threat intensity in controls, with higher levels of lever pressing for rewards during the tone (i.e. ‘riskier’ behavior) in mice trained using lower intensity foot shock. This suggests that KOs are more sensitive to threats and suppress reward seeking for lower-intensity shocks, while WTs balance reward/threat conflict based on shock magnitude. Critically, KOs were well-trained in reward acquisition prior to PMA conditioning, reaching press rates comparable to WTs, ruling out the possibility of deficits in baseline reward seeking explaining our results. Consistent with this finding, patients with OCD are generally more risk averse [(Frost & Steketee, 1994), but see (Luigjes et al., 2016)] and risk aversion is associated with reduced neural activity to rewards in cortico-striatal regions (Admon et al., 2012; Figee et al., 2011). This risk aversion and avoidance behavior is thought to significantly contribute to the reduced quality of life seen in OCD by interfering with normal daily activities (Huppert et al., 2009). These findings highlight the value of testing reward conflict in the PMA task, which we have now optimized for mice, building on a recent publication that developed a platform avoidance task without reward conflict in mice (Halcomb et al., 2024).

Following PMA acquisition and expression, KOs and WTs were trained on ERP to determine if KOs would have impaired extinction learning, consistent with phenotypes observed in fear conditioning paradigms (Kajs et al., 2022). In line with our predictions, KOs were slower to reengage in reward-seeking during tone periods, suggesting delayed extinction. Specifically, conditioning with stronger foot shocks (0.4 mA) produced persistent suppression of lever pressing in KOs, as evidenced by genotype differences in suppression to the first tone in the extinction test session. In contrast, the similar reduction of platform avoidance observed in KOs and controls following ERP (when comparing the end of conditioning and the extinction test sessions), indicates that conditioned avoidance responses were successfully extinguished in both groups. Curiously, conditioning with milder foot shocks produced lower platform time following ERP in KOs compared to WTs, counter to our predictions that KOs might show persistent avoidance similar to the effect of avoidance overtraining in rats (Martinez-Rivera et al., 2020) and avoidance devaluation in patients with OCD (Gillan et al., 2014).

To identify cortical and striatal regions that may be involved in regulating conflict-based avoidance and differentially active in *Sapap3*-KOs, we quantified c-Fos expression following the extinction test session in mice trained on the low intensity 0.24 mA shock, who displayed the most pronounced genotype differences in lever press suppression during conditioning. We found region-specific genotype differences in c-Fos expression, providing preliminary evidence for decreased mOFC c-Fos in *Sapap3*-KOs after avoidance reinstatement following ERP. The mOFC has been found to be hyperactive during initial avoidance acquisition in OCD patients (Gillan et al., 2015), whereas a role in later expression or persistence of avoidance has not been identified. It will be important to address this discrepancy in future investigations examining mOFC neural activity *in vivo* across avoidance acquisition, extinction, and extinction test. In addition to genotype differences, we identified associations between greater c-Fos density in the mOFC and increased lever suppression following the first tone after ERP training, as well as a trend towards IL c-Fos positively correlating with lever suppression. In line with our findings, studies in human participants have found positive associations between successful fear extinction and vmPFC activity (encompassing mOFC) (Milad & Rauch, 2007), as well as associations between increased vmPFC – amygdala functional connectivity and learned threat controllability (i.e. avoidance) (Boeke et al., 2017; Wanke & Schwabe, 2020). In addition, preclinical studies have identified an important role of mOFC and IL in regulating freezing behaviors in fear conditioning and fear extinction (Diehl et al., 2019; Moscarello & LeDoux, 2013; Rodriguez-Romaguera et al., 2015), although more work is needed to determine the role of these and other structures in flexible avoidance behavior under motivational conflict (Manning, Bradfield, et al., 2021; Martinez-Rivera et al., 2023; Rodriguez-Romaguera et al., 2016). mOFC also plays an important role in risk/reward decision making, with mOFC inactivation biasing risky choices in probabilistic discounting for rewards (Stopper et al., 2014); this may be relevant to our findings of lower suppression of lever pressing in animals with reduced mOFC c-Fos. Taken together, our findings and those in the literature support the idea that mOFC and IL are capable of influencing distinct aspects of post ERP behavior in the PMA task and OCD relevant models. This idea warrants further investigation with more precise methods such as *in vivo* recordings or temporally specific optogenetics.

Importantly, the scope of our study was limited to assessing active avoidance (i.e. time spent on the platform) and did not analyze differences in freezing behaviors. Previous studies in the *Sapap3*-KO model have found heightened freezing responses during fear conditioning and impaired extinction of freezing responses (Kajs et al., 2022). Deletion of *Sapap3* may produce a phenotype with heightened passive, rather than active, avoidance behaviors, due to an inability to flexibly switch defensive strategies coupled with greater sensitivity to contextual and cued threats to elicit fear responses. As active avoidance requires a process of fear reduction to switch from passive (i.e. freezing) to active (i.e. flight, escape) behaviors (LeDoux et al., 2017; Mowrer, 1956), quantifying freezing throughout avoidance acquisition and extinction is critical for determining if the current observations arise from increased passive avoidance, or from differences in reward valuation in negative contexts. In addition, only male mice were used in these studies, so generalizability of these results to females must be tested in future studies [Note: prior work from our lab and others has found no evidence for sex differences across any phenotype tested in *Sapap3*-KOs (Manning et al., 2019; Manning, Wang, et al., 2021; Pinhal et al., 2018)]. Regardless, both preclinical and neuroimaging studies of fear conditioning and active avoidance suggest that medial prefrontal cortices are critical for gating the selection of passive and active defensive responses (Diehl et al., 2019; Diehl et al., 2018; Moscarello & LeDoux, 2013; Sotres-Bayon et al., 2006). In summary, our work provides further rationale for monitoring and parsing the role of infralimbic and medial orbital cortices in regulating PMA and active avoidance, acquisition, and extinction, and identifies the mOFC as a target for alterations in conflict-based avoidance in a transgenic model relevant to OCD.

## Supplementary figures

**Supplemental Figure 1.**
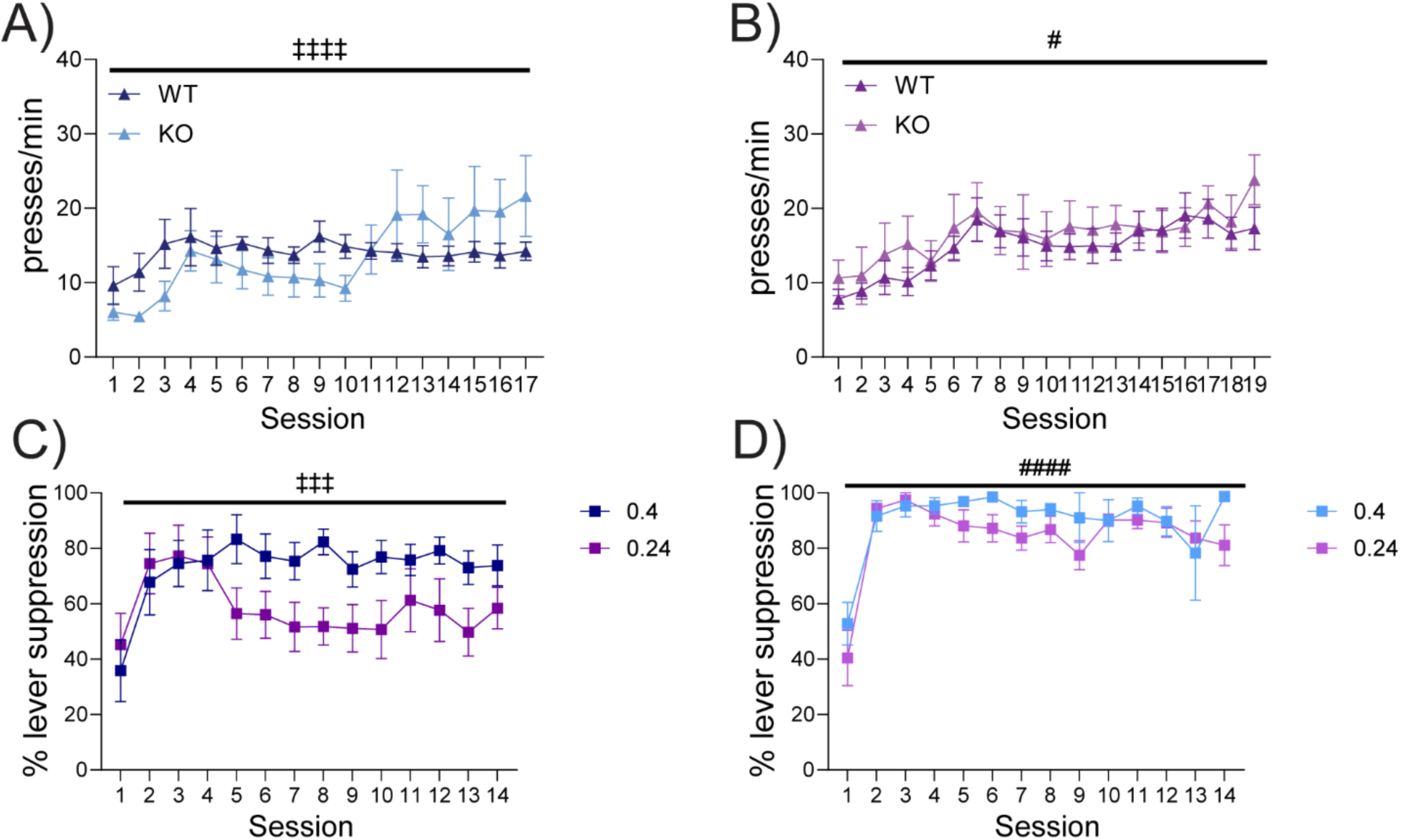
Trained mice initially seek rewards at similar rates, but suppression is genotype specific. A, B) Reward training prior to avoidance sessions produces similar performance in WT and KOs in 0.4 (A) and 0.24 (B) cohorts. C) Time-dependent differences in lever suppression emerges in WTs trained with different shock intensities (0.4: n=7; 0.24: n= 7). D) KOs exhibit high levels of lever suppression with platform avoidance conditioning, independent of shock intensity (0.4: n=5; 0.24: n=7). # - Time effect; ‡- Interaction (time x genotype or time x shock intensity); ‡ - p<0.05; ‡‡‡ - p<0.001; ####,‡‡‡‡ -p<0.0001.

**Supplemental Figure 2:**
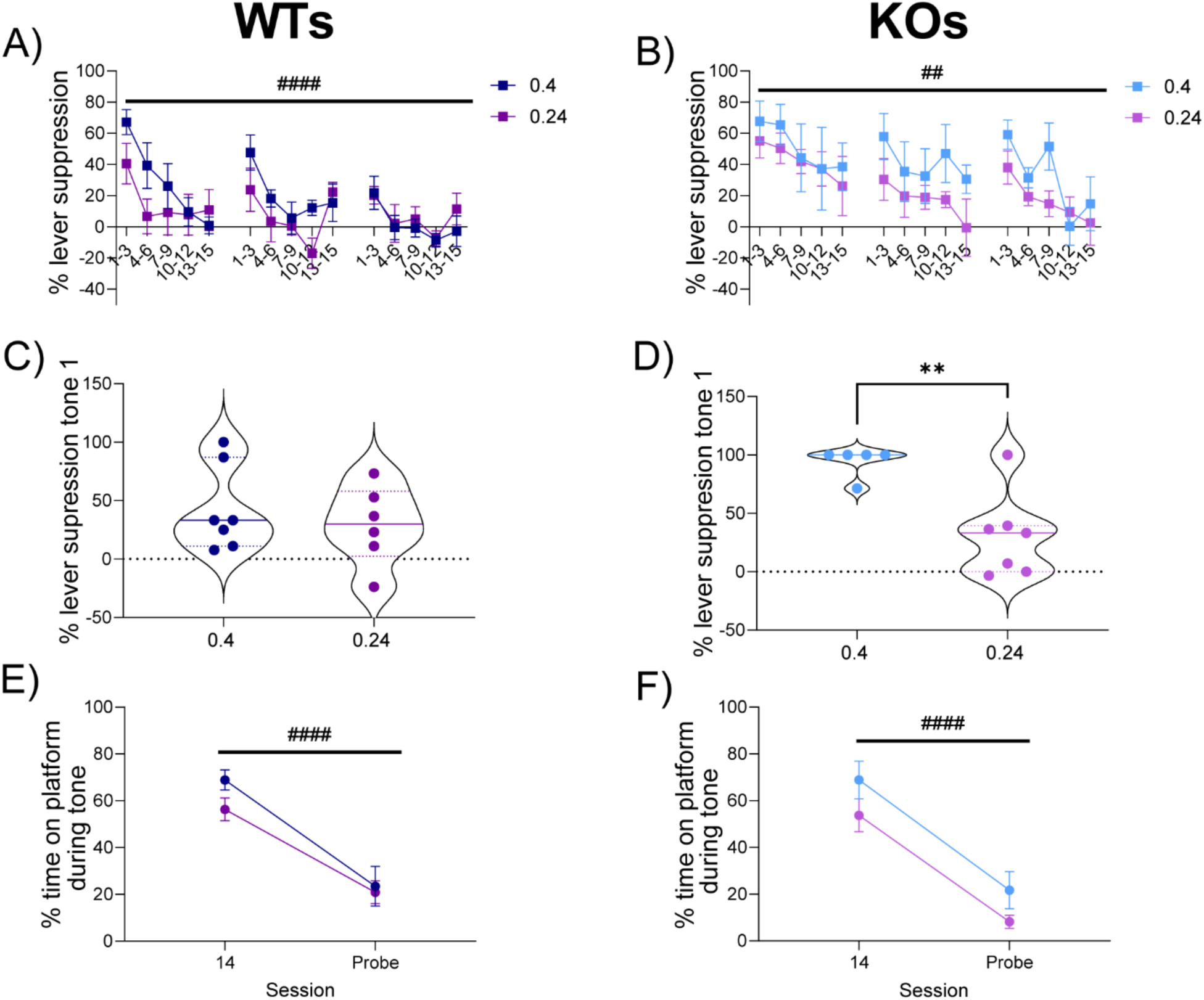
No differences in extinction of lever suppression and avoidance by shock intensity. A, B) WTs (A) and KOs (B) extinguish lever suppression (i.e. resume reward seeking) over ERP at similar rates, regardless of shock intensity during conditioning. C) Lever suppression at the extinction test to the first tone is similar in WT cohorts. D) Conditioning with higher intensity foot shock (0.4 mA) produces reinstatement of lever suppression in KOs compared to milder foot shock at the extinction test. E, F) WTs (E) and KOs (F) have no differences in platform avoidance reinstatement based on shock intensity during conditioning. WTs (0.4 mA: n=7, 0.24 mA: n=7); KOs (0.4 mA: n=5, 0.24 mA: n=7). #-main effect of time, *-main effect of shock intensity. ##, **- p<0.01; #### - p<0.0001.

